# 3DMolMS: Prediction of Tandem Mass Spectra from Three Dimensional Molecular Conformations

**DOI:** 10.1101/2023.03.15.532823

**Authors:** Yuhui Hong, Sujun Li, Christopher J. Welch, Shane Tichy, Yuzhen Ye, Haixu Tang

## Abstract

**Motivation:** Tandem mass spectrometry is an essential technology for characterizing chemical compounds at high sensitivity and throughput, and is commonly adopted in many fields. However, computational methods for automated compound identification from their MS/MS spectra are still limited, especially for novel compounds that have not been previously characterized. In recent years, *in silico* methods were proposed to predict the MS/MS spectra of compounds, which can then be used to expand the reference spectral libraries for compound identification. However, these methods did not consider the compounds’ three-dimensional (3D) conformations, and thus neglected critical structural information.

**Results:** We present the **3D Mol**ecular Network for **M**ass **S**pectra Prediction (3DMolMS), a deep neural network model to predict the MS/MS spectra of compounds from their 3D conformations. We evaluated the model on the experimental spectra collected in several spectral libraries. The results showed that 3DMolMS predicted the spectra with the average cosine similarity of 0.687 and 0.475 with the experimental MS/MS spectra acquired in positive and negative ion modes, respectively. Furthermore, 3DMolMS model can be generalized to the prediction of MS/MS spectra acquired by different labs on different instruments through minor fine-tuning on a small set of spectra. Finally, we demonstrate that the *molecular representation* learned by 3DMolMS from MS/MS spectra prediction can be adapted to enhance the prediction of chemical properties such as the elution time (ET) in the liquid chromatography and the Collisional Cross Section (CCS) measured by ion mobility spectrometry (IMS), both of which are often used to improve compound identification.

**Supplementary information:** The codes of 3DMolMS is available at https://github.com/JosieHong/3DMolMS and the web service is at https://spectrumprediction.gnps2.org.

## 1 Introduction

Because of its high sensitivity and throughput, mass spectrometry (MS) coupled with gas chromatography (GC) or liquid chromatography (LC) has long been adopted for the characterization and structural elucidation of chemical compounds (Vinaixa *et al.*, 2016). More recently, liquid chromatography tandem mass spectrometry (LC-MS/MS), which detects the fragment ions resulting from the high energy collision of compounds in a collision cell has become an essential technology for identifying and quantifying chemical compounds in complex samples with applications in multiple domain areas such as metabolomics (Alseekh *et al.*, 2021), natural product discovery (Hoffmann *et al.*, 2014), and environmental science (Richardson, 2008). For instance, metabolomics aims to identify and quantify metabolites present in tissues and body fluids, leading to the discovery of molecular biomarkers associated with diseases and clinical conditions (A Nagana Gowda and Raftery, 2013). Specifically, in untargeted metabolomics, LC-MS/MS is used to acquire thousands of MS/MS spectra in a single sample, from which metabolites are to be characterized (Xiao *et al.*, 2012). Many MS-based metabolite identification systems exploited the spectra searching against a reference spectral library (RSL) consisting of the MS/MS spectra of previously identified chemical compounds (Stein, 2012). In practice, however, the current spectral libraries (e.g., NIST20 (Yang *et al.*, 2017), HMDB (Wishart *et al.*, 2022), MassBank (Horai *et al.*, 2010) and GNPS (Wang *et al.*, 2016)) only contain the experimental MS/MS spectra of a limited number of compounds (e.g., the NIST20 and GNPS spectral libraries contain the MS/MS spectra of 29,890 and 18,163 compounds, respectively, while the other libraries are even smaller), and thus a majority (i.e., over 80%) of MS/MS spectra in metabolomic experiments remain un-identified by the spectral library searching methods (Griss *et al.*, 2016). Notably, compound identification remains a big obstacle in the other applications of LC-MS/MS such as in environmental chemistry (Richardson, 2008) and natural product discovery (Hoffmann *et al.*, 2014), in which the fraction of unknown compounds in a target sample is even greater.

To address this problem, computational methods have been developed for compound identification from their MS/MS spectra (Blaženović *et al.*, 2018). Among them, one approach aims at the *in silico* prediction of MS/MS spectra from compounds, and then uses the predicted spectra of potential small molecules in the entire chemical space (e.g. those in the PubChem database) for the spectral library searching. Many of the spectral prediction algorithms adopted a rule-based approach, in which empirical and/or computer-learned rules are devised and combined to derive the fragment ions and their intensities, e.g., those implemented in LipidBlast (Kind *et al.*, 2013), MetFrag (Ruttkies *et al.*, 2016), MIDAS (Wang *et al.*, 2014), and CFM-ID (Wang *et al.*, 2021). The state-of-the-art rule-based method for predicting MS/MS spectra of compounds is CFM-ID 4.0 (Wang *et al.*, 2021). Given the chemical structure of a compound, CFM-ID first computes a feature vector of potential fragments and then uses empirical fragmentation rules to compute the putative chemical bond cleavages and the charges of these fragments. Finally, an MS/MS spectrum is predicted by combining the empirical and computer-learned (by using machine learning algorithms) fragmentation rules.

Recently, with the cumulative MS/MS spectra of small molecules enabled by the efforts of several large-scale spectral libraries (e.g., GNPS (Wang *et al.*, 2016)), it has become plausible to train deep learning models for predicting MS/MS spectra of chemical compounds. In fact, deep learning models have been successfully adopted to predict the MS/MS spectra of peptides, not only for the prediction of the fragment ions’ intensities (Zhou *et al.*, 2017; Zeng *et al.*, 2019; Tarn and Zeng, 2021; Gessulat *et al.*, 2019; Tiwary *et al.*, 2019; Yang *et al.*, 2020) but also for the prediction of full MS/MS spectra from scratch (Liu *et al.*, 2020), which demonstrated the power of deep neural networks (DNNs) for learning complex rules and patterns from massive training data. Despite the success of DNNs in peptide MS/MS spectra prediction, their application to compound spectra prediction is limited, mainly because unlike peptides that can be represented as linear sequences, small molecules are represented as molecular graphs, and thus the sequence-to-sequence deep learning models cannot be directly exploited. On the other hand, machine learning methods have been widely used to predict the chemical properties of compounds. For these prediction tasks, an essential issue is how to represent chemical compounds as the input to the DNNs. A rule-based method takes as input the stereochemical information of the compounds by encoding them into *fingerprints*, like Extended-Connectivity Fingerprints (ECFP) (Rogers and Hahn, 2010) and MinHashed atom-pair fingerprints up to a diameter of four bonds (MAP4) (Capecchi *et al.*, 2020). However, rule-based encoding limits the capabilities of deep learning models, which are capable of learning new rules directly from training data. The current DNNs for learning compound representations fall into three categories: SMILES-based methods, graph-based methods, and point-based methods. A SMILES-based convolutional neural network (CNN) (Hirohara *et al.*, 2018) encodes the SMILES strings by one-hot coding, while CHEM-BERT (Kim *et al.*, 2021a) encodes SMILES strings as embedding vectors, both of which do not contain any structural information of compounds. The graph-based methods encode a compound as a molecular graph, in which the nodes represent atoms, and the edges represent chemical bonds, and then exploit the molecular graph as the input to various graph neural networks (GNNs), e.g., the Graph Convolutional Network (GCN) (Wang *et al.*, 2022) or the Message Passing Neural Network (MPNN) (Gasteiger *et al.*, 2020; Klicpera *et al.*, 2020). However, the bond types are restricted in graph-based methods; for instance, intra-molecular hydrogen bonds are not directly modeled. The point-based methods exploit the atomistic systems (chemical bonds are not taken into account by these methods) and encode the atoms as points in DNNs, e.g. in SchNet (Schütt *et al.*, 2018) and the Polarizable atom interaction neural network (PAINN) (Schütt *et al.*, 2021). In SchNet, the continuous-filter convolutional layers are designed to model the atoms, but only the distances among atoms are considered. PAINN makes use of not only the distances between atoms but also the angles between bonds. To the best of our knowledge, NEIMS (Wei *et al.*, 2019) and MassFormer (Young *et al.*, 2021) are the only existing neural network models for the compound MS/MS spectra prediction, which encode the small molecules as ECFP and molecular graphs, respectively, and then exploit the multilayer perceptron (MLP) neural network and Graph Transformers (Yun *et al.*, 2019) for spectra prediction, respectively.

In this paper, we present an efficient deep neural network, 3DMolMS, based on an elemental operation of three-dimensional (3D) molecular convolution on the 3D molecular conformations of compounds to predict their MS/MS spectra. In 3DMolMS, the 3D conformations are represented as *point-sets* (Qi *et al.*, 2017; Manessi *et al.*, 2020), which is a commonly used representation in computer vision to simulate the objects in the real world. The molecular point-sets encode the accurate 3D coordinates and attributes of the atoms, while the chemical bonds are represented in *neighboring vectors.* When trained and tested on the compounds’ MS/MS spectra collected from the Agilent Personal Compound Database and Library (PCDL) and NIST20, 3DMolMS achieved higher accuracy than CFM-ID 4.0 (Wang *et al.*, 2021), NEIMS (Wei *et al.*, 2019) and the existing deep learning model MassFormer (Young *et al.*, 2021). Furthermore, 3DMolMS achieves satisfactory performance when applied to the prediction of the spectra in the public spectral library MassBank of North America (MoNA) with minor fine-tuning, which indicates the 3DMolMS model can be generalized to the prediction of spectra acquired by different labs on different instruments. Finally, we showed the representation learned in spectra prediction can be adapted to enhance the prediction of chemical properties of compounds such as the elution time (ET) in liquid chromatography, and the collisional cross section (CCS) in the ion mobility spectrometry (IMS), both of which are often used as complementary measures in compound identification.

## 2 Materials and methods

### 2.1 MS/MS spectral libraries

To train and evaluate the deep learning model for MS/MS spectra prediction, we gathered about 70,000 spectra, all acquired using the Agilent quadrupole/time-of-flight (Q-TOF) MS instruments, from the Agilent Personal Compound Database and Library (PCDL) and the NIST20 spectral library (Yang *et al.*, 2017) (Table 1). We combined these two libraries for training purposes because 1) although PCDL contains the spectra from a large number (11, 239) of compounds, NIST20 provides somewhat complementary data: it collected the experimental MS/MS spectra from 2,492 compounds, among which only 532 (i.e., 21%) compounds are also collected by PCDL; 2) NIST20 collected multiple spectra from the same compound (on average, 11 spectra per compound), which can be exploited by the deep learning models to model the experimental variations among replicated MS/MS acquisitions; and 3) most (96%) Q-TOF MS/MS spectra in NIST20 were acquired by using Agilent Q-TOF instruments (we only kept these spectra in our training and testing data), and as a result, the MS/MS spectra of the compounds shared by both libraries are highly similar, with the average cosine similarities of 0.870 and 0.982 for the MS/MS spectra acquired in the positive and negative ion modes, respectively. Notably, we used the experimental MS/MS spectra of 90% randomly selected compounds in the combined library for the training, and the spectra of the remaining 10% compounds for testing to ensure the training and testing datasets are not overlapping with each other.

**Table 1:**
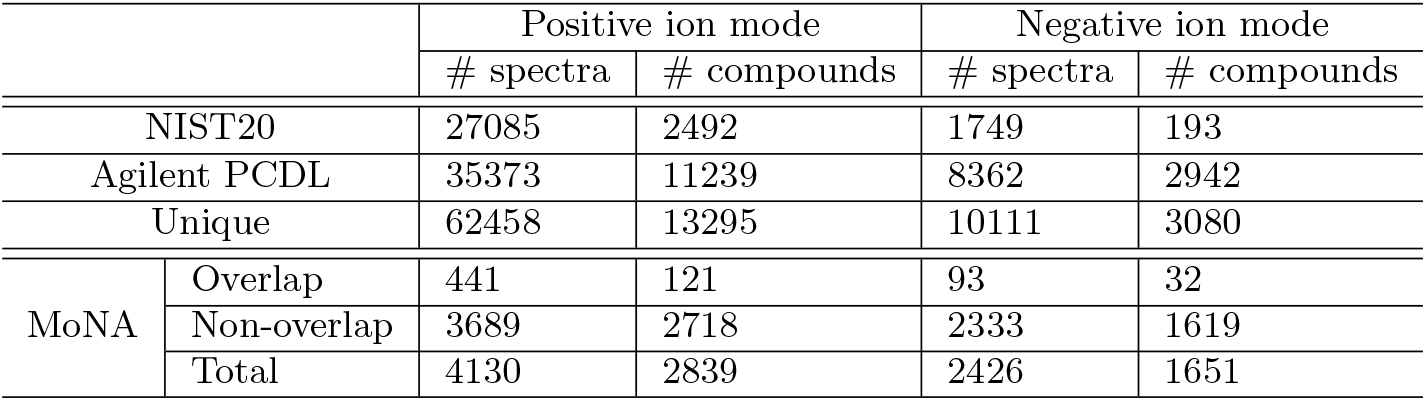
Total numbers of spectra and compounds used for training and testing the spectra prediction models. The compounds (and their MS/MS spectra) in the MoNA library that are also present in the PCDL or the NIST20 libraries are counted as “overlap”, and the remaining are counted as “non-overlap”.

|  |  | Positive ion mode |  | Negative ion mode |  |
| --- | --- | --- | --- | --- | --- |
|  |  | # spectra | # compounds | # spectra | # compounds |
| NIST20 |  | 27085 | 2492 | 1749 | 193 |
| Agilent PCDL |  | 35373 | 11239 | 8362 | 2942 |
| Unique |  | 62458 | 13295 | 10111 | 3080 |
| MoNA | Overlap | 441 | 121 | 93 | 32 |
|  | Non-overlap | 3689 | 2718 | 2333 | 1619 |
|  | Total | 4130 | 2839 | 2426 | 1651 |

We also selected about 6, 000 experimental Q-TOF MS/MS spectra from MassBank of North America (MoNA) (Horai *et al.*, 2010) as an independent test set. We note that the specific MS instrument used to acquire these MS/MS spectra were not recorded in the library: some of them may be acquired by using the Agilent instrument, while the others may be acquired by using instruments from other vendors. Furthermore, the specific collision energies to acquire the MoNA spectra were not recorded either. Hence, the similarity between the MS/MS spectra of the 121 compounds shared in MoNA and the combined PCDL/NIST20 library is relatively low, with the average cosine similarities of 0.655 and 0.804 for the MS/MS spectra of the positive and negative parent ions, respectively. We used the MoNA library as independent testing data to demonstrate the capability and limitation of the deep learning model for predicting the MS/MS spectra acquired by different MS instruments under different experimental conditions.

For all three libraries, the MS/MS spectra were first filtered with the following steps: (1) The missing isomeric SMILES are fixed by searching with the synonyms names in PubChem (Kim *et al.*, 2021b); (2) The mass spectra have less than 5 peaks are filtered out because they are typically unreliable. (3) The m/z range is limited in (0, 1500], because few spectra have m/z above 1500. (4) Only the molecules composite by the high-frequency atoms (C, H, O, N, F, S, Cl, P, B, I, Br) are retained; (5) Only the spectra with high-frequency precursor types ([M + H]^+^, [M – H]^-^, [M + Na]^+^, [M + H – H_2_O] ^+^, [M + 2H]^+^) are retained. The statistics of the datasets after filtration are summarized in Table 1.

### 2.2 Encoding 3D conformations of compounds

We used the ETKDG (Riniker and Landrum, 2015) algorithm implemented in the `AllChem.EmbedMolecule()` function of the RDkit library (Landrum *et al.*, 2020) to generate the 3D conformations (also referred to as the *conformers*) of each compound. The function takes as input the SMILES (i.e., simplified molecular-input line-entry system) string of each compound, and outputs the 3D conformer in the `sdf.gz` format, which contains the *x, y, z*-coordinates of each atom in the compound. The conformer of a compound is then encoded into a fixed number of *N* points of atoms (i.e., the *point set*), where *N* is the maximum number of atoms in the considered compounds (by default *N* = 300). When the number of atoms in a compound is smaller than *N*, the point set is padded to *N* points with the coordinates of the padded points set to be zeros. Each atomic point is encoded by the *x, y, z*-coordinates as well as a number of attributes of the atom, as summarized in Table 2; the atom attributes are generated by using RDKit.

**Table 2:** The point set encoding of a compound, in which each atom in the compound is encoded as a vector of 21 dimensions, representing the *x, y, z*-coordinates and other attributes of the atom.

| Index | Description |
| --- | --- |
| 0-2 | x, y, z coordinates |
| 3-14 | one-hot encoding of the atom type |
| 15 | number of immediate neighbors who are nonhydrogen atoms |
| 16 | valence minus the number of hydrogens |
| 17 | atomic mass |
| 18 | atomic charge |
| 19 | number of implicit hydrogens |
| 20 | is aromatic |
| 21 | is in a ring |

### 2.3 Encoding MS/MS spectra

We represent an experimental MS/MS spectrum as a 1D spectral vector, in which each dimension represents the total intensity of fragment ions in a bin of the fixed mass-to-charge ratio (m/z). Here, the total number of bins (i.e., the dimension of the spectral vector) is dependent on the mass resolution of the MS/MS spectrum and is set as an adjustable parameter in our model. By default, the resolution is set to be 0.2, resulting in the spectral vector of 7, 500 dimensions (within the m/z range between 0 and 1, 500 that covers almost all fragment ions observed in the MS/MS spectra of compounds).

In practice, the peak intensities are often transformed to reduce the impact of the most intensive peaks when computing the similarities between spectra, in particular, to separate the similarity distribution of replicated spectra of the same compound from the distribution between the spectra of different compounds. Based on the previous studies (e.g., for peptide spectra prediction (Liu *et al.*, 2020)), we adopted the square root transformation, i.e., the intensity of each peak is converted into its square root. We note that the logarithm transformation is also used in previous studies (Lam *et al.*, 2006), which can achieve a similar effect.

Finally, we consider the MS experimental conditions, including the precursor types and the collision energy as the metadata, which are concatenated to the embedded point set. The precursor types are encoded in one-hot codes, while the collision energies (recorded in eV in the PCDL and NIST20 libraries) are converted into the normalized collision energy (NCE) by using Eq. 1.

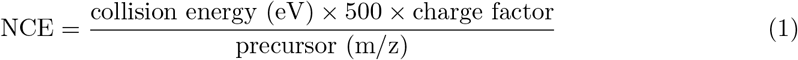

in which the charge factor is 0.9 for doubly charged molecules. As not all the spectra (about 78%) are labeled with collision energy in MoNA, the unlabeled ones are set to be 20 eV.

### 2.4 3DMolConv

Point-based DNNs are widely used in computer vision (Qi *et al.*, 2017; Manessi *et al.*, 2020; Schütt *et al.*, 2018, 2021). The main idea of these models is to apply a *symmetric* function to the elements in the point set to extract representation features of the global 3D structure. Formally, given a point set, *X* = {*x*_1_, *x*_2_,…,*x_n_*} with *x_i_* ∈ ℝ^*F*^, the representation of the point set can be obtained by applying a series of the symmetric functionh:

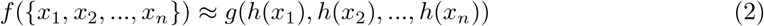

where *f* : 2^R^^F^ → ℝ is the representation function on the input point set, which is summarized from the elemental operation *h* : ℝ^*F*^ → ℝ^*K*^ on each element *x_i_* in the point set through an aggregated function 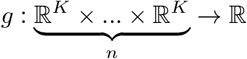.

Here, we adopt the max-pooling function for *g*, and the sequential convolutional layers as *h*. Specifically, for each layer *l*, we design the symmetric function (referred to as the elemental operation *3DMolConv*) as:

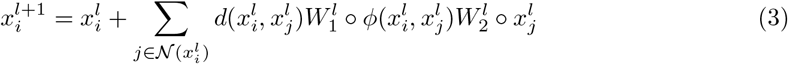

where ○ represents the element-wised multiplication, the 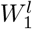 represents the filter on the distance, 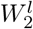 represents the filter on the direction, and 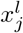 represents one point (i.e., *x_j_*) of the *k*-nearest neighbors of the point *x_i_*. Here, the distance between two points *x_i_* and *x_j_* is computed as:

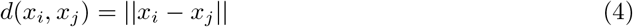

and the angle between the point vectors *x_i_* and *x_j_* encodes the information related to either the bond angle or the non-bond angles of the edge 〈*x_i_, x_j_*〉:

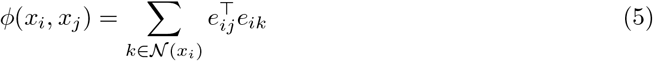

where *e_ij_* denotes the vector representation related to the edge *e_ij_* between *x_i_* and *x_j_*:

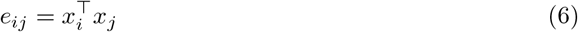

We show in Fig. 1 that the angles (or the inner product) between two vectors can encode either the bond or the non-bone angles between the two points, which have the equivalent effect as the encoding of torsion angles (Adams *et al.*, 2021) to represent compounds. Compared to the vector subtraction (*e_ij_* = *x_i_* – *x_j_*) used by DGCNN (Manessi *et al.*, 2020), the inner product representation retained more structural information about the points. In addition, we prove that 3DMolConv satisfies the permutation invariance and the SE(3) (i.e., the Euclidean group subject two 3D displacement motions such as translations and rotations) invariance in Section S1.

**Figure 1:**
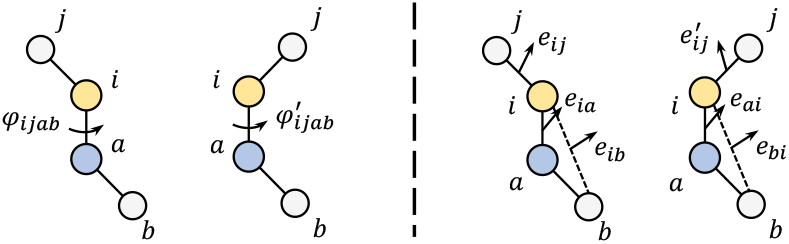
A schematic illustration of message passing to the point (atom) *i* from one of its nearest neighbors (atom) *j* in two point-sets (representing two different chemical structures) using the torsion angle representation (left) and the representation of the angles between these two vectors (right), respectively, on a compound with *N* = 4 points (and the number of nearest neighbors of *k* = 3). Obviously, the angle representation involves only pairs of points, while can distinguish these two different chemical structures (i.e., 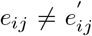). As a result, this representation has the equivalent effect as the encoding of torsion angles (i.e., 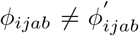) that involves four points (atoms).

### 2.5 Model architecture of 3DMolMS

Based on the elemental operation 3DMolConv, we constructed the 3DMolMS model as illustrated in Fig. 2. The input of the network is the encoding vectors of the *x, y, z*-coordinates and attributes of all atoms in a compound, represented in a matrix. The input matrix is then processed by an encoder consisting of six 3DMolConv-based hidden layers with the dimensions of 64, 64, 128, 256, 512, and 1, 024, respectively. All molecular representation vectors from the hidden layers in the encoder are concatenated together with the metadata to form the final latent representation vector in 2, 048 dimensions. Notably, the *x, y, z*-coordinates are not directly encoded after the first layer in the encoder to ensure the rotation invariance of the model representation, as explained in Section S1. The latent representation vector from the encoder is then fed into a spectrum decoder consisting of five fully connected (FC) layers with residual blocks to output the predicted spectrum. All hidden layers in the decoder have the same dimension of 2,048 (see Fig. S1). After the predicted MS/MS spectrum is output by 3DMolMS, we eliminate all the peaks whose intensities are smaller than 0.001 (i.e., 0.1% of the total intensity of the spectrum), and then convert the intensities of the remaining peaks into their square in the final predicted spectrum.

**Figure 2:**
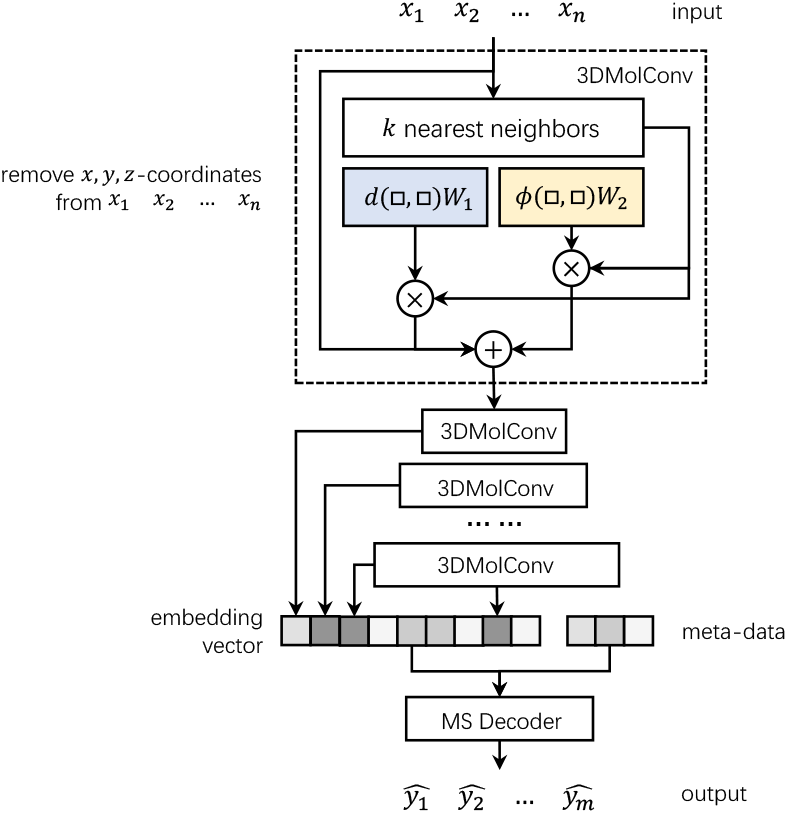
The architecture of the encoder used by 3DMolMS, which consists of six 3DMolConv-based hidden layers. All molecular representation vectors from these hidden layers are concatenated together with the metadata to form the final latent representation vector in 2, 048 dimensions. The latent vector is then converted by a decoder (consisting of five fully connected layers) into the final predicted spectral vector.

### 2.6 Transfer representation learning

Through 3DMolMS, each compound is embedded into a 2048 dimensional latent vector by the encoder, which contains sufficient information to predict the MS/MS spectra of the compound. We hypothesize that this molecular representation learned from MS/MS spectra prediction captures essential structural information about the compounds, which can be transferred to the relevant prediction tasks, such as the prediction of the chemical properties of compounds. Here, as a proof of concept, we demonstrate this transfer learning approach indeed improves the prediction of the elution time and the collisional Cross Section (CCS) (Plante *et al.*, 2019; Zhou *et al.*, 2020) and elution time (ET) (Domingo-Almenara *et al.*, 2019) of compounds. Specifically, the weights in the encoder of the pretrained spectra prediction model are directly used to generate the representation vector of each compound, which is then fed into another decoder to predict the respective property such as CCS or ET. When trained for a prediction task, the representation learning is fine-tuned by the respective training dataset of the task, while the decoder is trained independently. In this paper, we used the same decoder architecture for the prediction of CCS and ET, which includes five layers of fully connected (FC) layers.

## 3 Results

### 3.1 Performance of predicting MS/MS spectra acquired by using Agilent Q-TOF instruments

We implemented 3DMolMS in the PyTorch framework (Paszke *et al.*, 2017, 2019). It takes about 0.13 second to predict an MS/MS spectrum from a single input compound, while the additional 0.22 second is needed for the computation of the compound’s 3D structure using RDkit (Section S2). In comparison, it takes around 0.03, 0.05, and 24.1 seconds for NEIMS, MassFormer, and CFM-ID 4.0 to predict the MS/MS spectrum of a compound.

We compare our model against the existing mass spectra prediction methods, including CFM-ID 4.0 (Wang *et al.*, 2021), NEIMS (Wei *et al.*, 2019) and MassFormer(Young *et al.*, 2021). NEIMS and MassFormer are retrained by using our training set for a direct comparison. Because the training codes of CFM-ID are not publicly available and the precursor types supported by the software are different from ours, we selected the 3, 482 and 770 spectra in the testing set with the adduct ions of [M + H]^+^ and [M – H]^-^, respectively, for the evaluation. In addition, we adapted three baseline point-based deep learning methods for MS/MS spectra prediction, including SchNet (Schütt *et al.*, 2018) developed for molecular representation learning, and Point-Net (Qi *et al.*, 2017), and DGCNN (Manessi *et al.*, 2020) developed for computer vision tasks. All these models are used to embed the same input point set into a latent representation vector, which is combined with the same metadata and then converted by the same decoder into the final predicted MS/MS spectrum. Cosine distance is used as the loss function for training all the DNNs. The detailed settings of the comparative experiment are described in Section S3.

As shown in Fig. 3, 3DMolMS predicted the MS/MS spectra that are highly similar to the experimental spectra, with the cosine similarity of 0.687(±0.362) and 0.475(±0.378) for the adduct ions of [M + H]^+^ and [M – H]^-^, respectively. The performances of all point-based DNN models (3DMolMS, SchNet, PointNet, and DGCNN) are better than existing spectra prediction algorithms (MassFormer, NEIMS, and CFM-ID), while 3DMolMS outperforms the baseline point-based DNN models. It is worth noting that the performance of all deep learning methods (including 3DMolMS) performs better on the spectra acquired under the positive ion mode than those under the negative ion mode, which is due to the fact that the training set of the positive ion mode is substantially larger than that of the negative ion mode (Table 1). As shown in Fig 4, the predicted accuracy increased significantly with more training samples, and thus there is still plenty of room for further improvement of these models including 3DMolMS. Note that CFM-ID is designed for all Electrospray ionization (ESI) spectra. However, the spectra from different instruments may vary substantially. Hence, the predicted spectra by CFM-ID may not represent the best prediction for the experimental spectra of the same collision energy. We count the maximum cosine similarity between the experimental spectra and the three predicted spectra under different collision energies (10 eV, 20 eV, and 40 eV, respectively) in our statistics for CFM-ID for a fair comparison.

**Figure 3:**
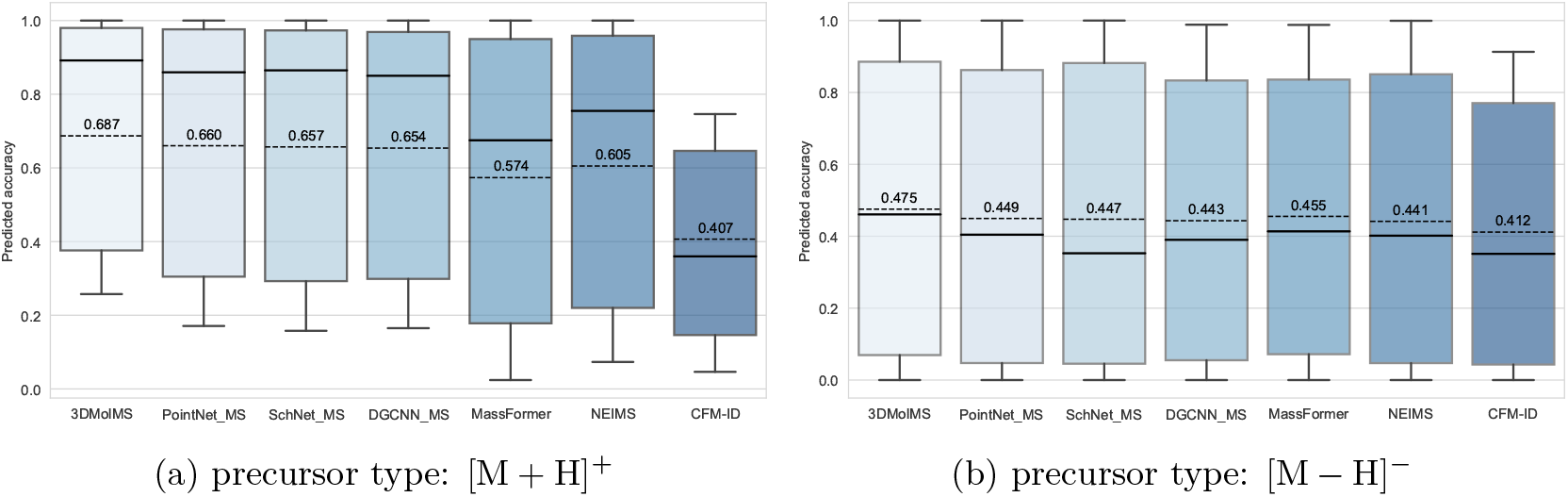
The boxplot of the cosine similarities between predicted and experimental MS/MS spectra acquired by using the Agilent Q-TOF instrument, in comparison with the similarities between the replicated experimental spectra from the same compounds. 3DMolMS are compared against the other point-based DNN models (PointNet, DGCNN, and SchNet) modified to MS prediction as well as the existing MS prediction models (MassFormer, NEIMS, and CFM-ID).

**Figure 4:**
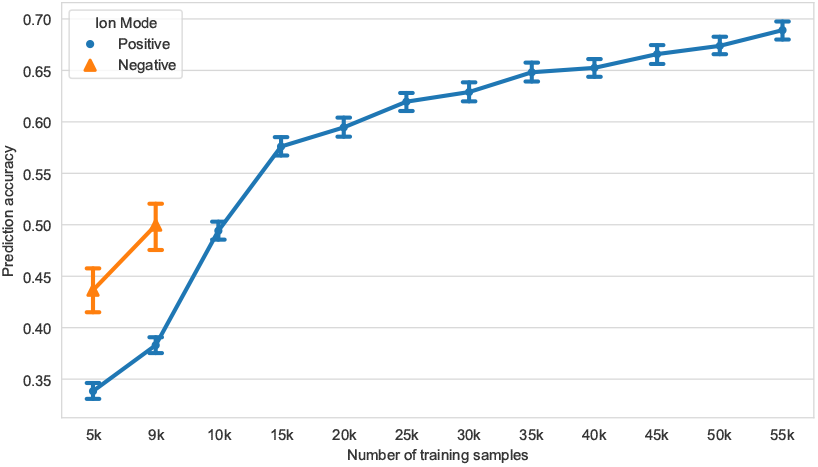
Prediction accuracy (measured by the similarity between the predicted and experimental spectra on testing data; y-axis) increases with more training data (x-axis).

Fig. 5 shows two examples of predicted MS/MS spectra by 3DMolMS, one for the positive ion mode and the other for the negative ion mode. In these cases, the predicted MS/MS spectra cover most of the fragment ion peaks, which indicates that the complex patterns of the 3D chemical structures of compounds associated with the fragmentation mechanisms can be learned by 3DMolMS, even though the fragmentation rules of compounds in tandem mass spectrometry remain unknown.

**Figure 5:**
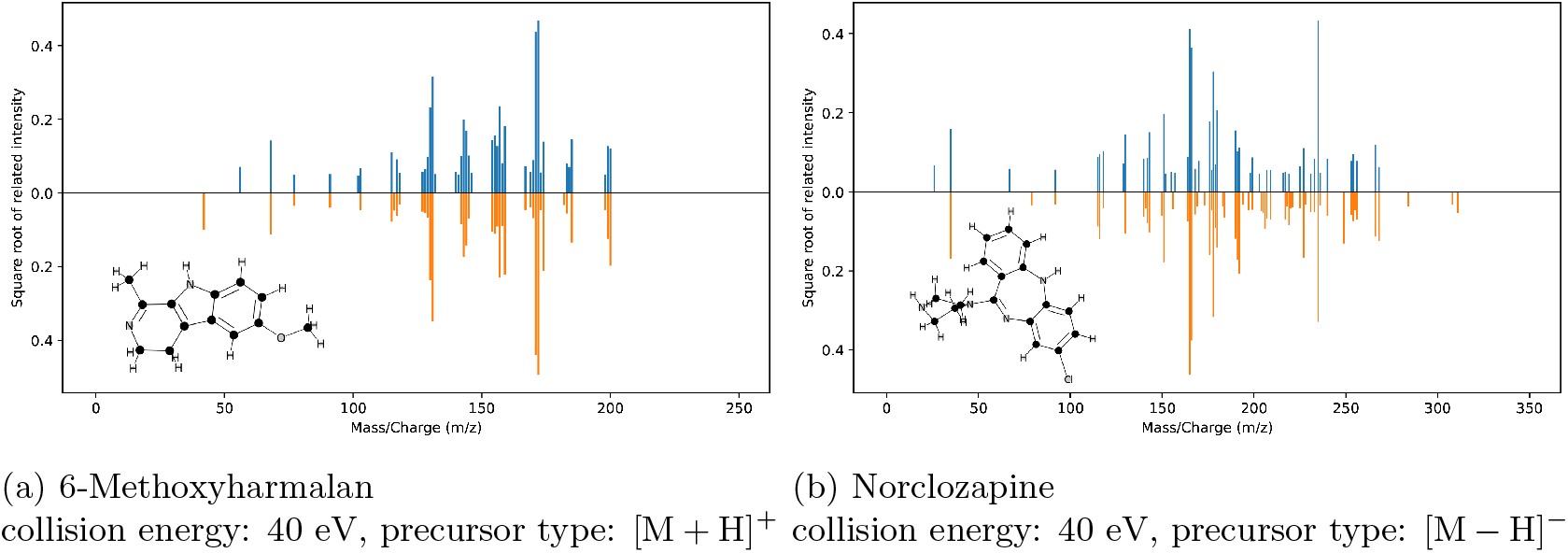
The examples of the predicted MS/MS spectra in comparison with the corresponding experimental spectrum of two compounds acquired in the positive and negative ion modes, respectively. The predicted spectra are labeled in orange (lower) and the experimental spectra are labeled in blue (upper).

As described in the Methods section, the collision energy is used as metadata in the 3DMolMS model, and thus 3DMolMS can predict the MS/MS spectra resulting from the fragmentation of the same compound under different collision energy. As shown in Table 3, the spectra predicted by 3DMolMS under specific collision energy are more similar to the experimental spectra of the same collision energy than those of different collision energy, which indicates the predicted spectra by 3DMolMS capture the distinctive fragmentation patterns under different collision energy. We note that here we consider two collision energy levels (20eV and 40eV) as examples to demonstrate the performance of spectra prediction under different collision energies because PCDL contains the MS/MS spectra of a sufficiently large number (1, 381 and 235 for positive and negative ion modes, respectively) of compounds under both energy levels in PCDL. The detailed distribution of the MS/MS spectra under different collision energies is shown in Fig. S2. We also observed that, for both ion modes, the prediction accuracy of 3DMolMS is higher on the spectra under 40eV than those under 20eV, perhaps due to on average more fragment ions being observed in the MS/MS spectra under higher collision energies, implying that the MS/MS spectra under higher collision energy may be more informative for compound identification.

**Table 3:** <u>First in the parentheses</u>: the average cosine similarities between the experimental and 3DMolMS-predicted MS/MS spectra; <u>Second in the parentheses</u>: the average cosine similarity between experimental spectra (in PCDL) under different collision energies.

|  | Positive ion mode |  | Negative ion mode |  |
| --- | --- | --- | --- | --- |
|  | 20 eV | 40 eV | 20 eV | 40 eV |
| 20 eV | (0.649, -) | (0.567, 0.600) | (0.467, -) | (0.385, 0.454) |
| 40 eV | (0.474, 0.600) | (0.758, -) | (0.257, 0.454) | (0.616, -) |

We then investigate the prediction performance of 3DMolMS across different types of compounds. As expected, we observed that the prediction performance of 3DMolMS on a compound (in the testing set) is correlated with its chemical similarity with the compounds in the training set. Here, we used the Bulk Tanimoto similarity (based on the chemical fingerprints) (Tanimoto, 1958) to measure the structural similarity between two compounds, and for each compound in the testing set, we measured its maximum similarity with any compound in the training set. As shown in Fig. S3, in general, the prediction performance increases when more similar compounds are present in the training set. It is not surprising that the model performs relatively better on the compounds with similar chemical structures because their similar compounds have been seen in the training set and thus the model has learned relevant fragmentation rules and patterns related to such compounds. This result also indicates the performance of 3DMolMS will be further enhanced if we expand the training set to cover the compounds with higher structural diversity. The performances of 3DMolMS using different diversity training samples in Fig. S4 agree with this hypothesis.

### 3.2 Extension of spectra prediction for other Q-TOF instruments

Next, we attempted to evaluate the performance of 3DMolMS on the prediction of MS/MS spectra acquired using the non-Agilent Q-TOF instruments (e.g., from the vendors of Bruker and Waters) collected in the MoNA spectral library. We first excluded about 150 compounds in the MoNA library that are also present in the PCDL or NIST20 libraries, and then randomly split the remaining compounds and their spectra into two subsets: one subset with 50% compounds for fine-tuning, and the other subset with 50% compounds for testing. As shown in Fig. 6, when we trained a 3DMolMS model from scratch on the fine-tuning subset, the predicted QTOF MS/MS spectra on the testing subset of the MoNA library achieved the cosine similarities of 0.445(±0.379) and 0.236(±0.300) for the positive and negative ion modes, respectively. On the other hand, when we fine-tuned the 3DMolMS model trained on the combined PCDL/NIST20 libraries using the fine-tuning subset, the prediction accuracies were significantly enhanced: the predicted spectra achieved the cosine similarities of 0.506(±0.316) and 0.430(±0.322) for the positive and negative ion modes spectra in the same testing data, respectively. For comparison, the spectra predicted by CFM-ID (following the same procedure described in Section 3.1) achieved the cosine similarities of 0.076(±0.127) and 0.291(±0.248) for the positive and negative ion modes spectra, respectively. The other spectra prediction methods (NEIMS and MassFormer) performed even more poorly, perhaps because of the small training dataset used here. Note that the spectra in the MoNA library are acquired from various Q-TOF instruments. As a result, the similarities between replicated spectra of the same compounds are not as high. As shown in Figure 6, the average cosine similarities between the spectra of the same compounds in the MoNA library are 0.655 and 0.804 for the spectra acquired in the positive and negative ion modes, respectively, which are much lower than the similarities (i.e., 0.870 and 0.982, respectively) in the combined PCDL/NIST20 libraries that were acquired by using the Agilent Q-TOF instrument. Hence, our results showed 3DMolMS may be extended to the prediction of MS/MS spectra acquired by different instruments after the fine-tuning on a small set of experimental spectra, even though the prediction accuracy might be relatively lower than the original model trained on the large training set. We also note the performance of 3DMolMS is better on the spectra in positive ion mode than on those in negative ion mode, because of the fewer training samples in negative ion mode in both the combined PCDL/NIST20 libraries used to train the original model and the MoNA library for fine-tuning.

**Figure 6:**
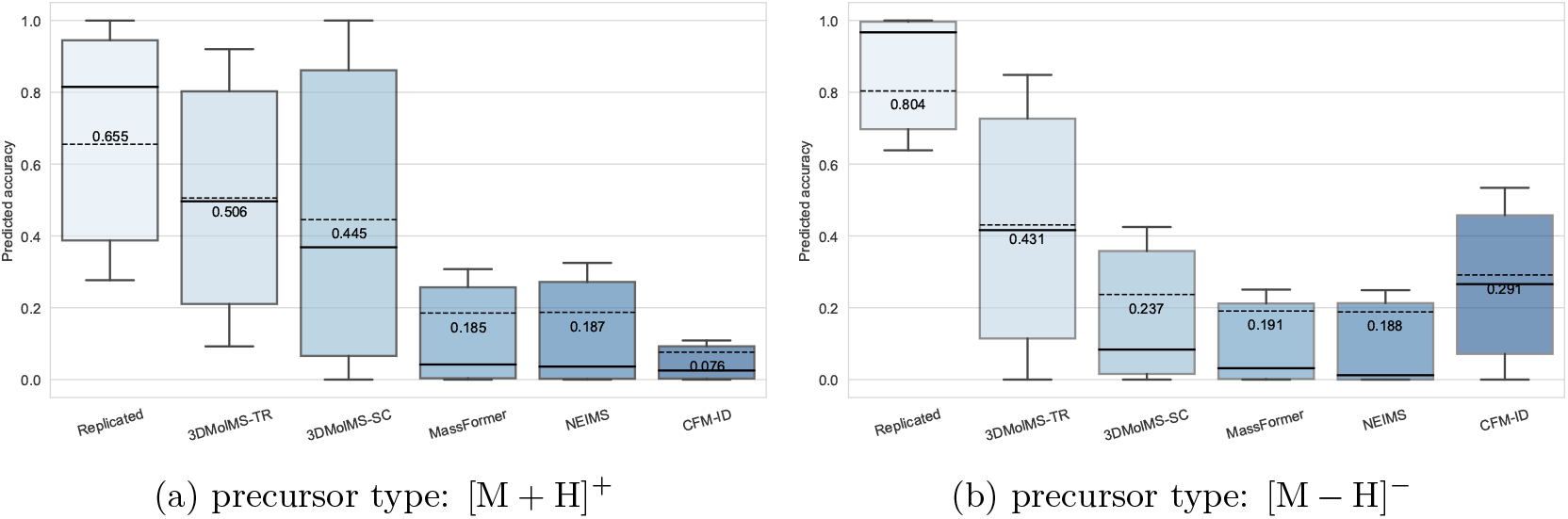
The cosine similarities between the predicted and the experimental MS/MS spectra in the MoNA library acquired by using various Q-TOF instruments, in which spectra prediction was performed by 3DMolMS, including the models fine-tuned (denoted as 3DMolMS-TR) or trained from scratch (denoted as 3DMolMS-SC), respectively, and existing spectra prediction models (MassFormer, NEIMS, and CFM-ID). The replicated similarities between the spectra of a small number of compounds shared by the combined PCDL/NIST20 libraries and the MoNA library are also shown here for comparison.

### 3.3 Transfer learning for the prediction of collision cross section and elution time

Finally, we investigate the transferability of the latent molecular representation learned from spectra prediction to the tasks of predicting chemical properties of compounds, specifically, their elution time (ET) in liquid chromatography (LC) and their collisional cross section (CCS) measured by ion mobility spectrometry (IMS). For these two regression tasks, we utilized the training datasets collected in previous studies to build customized machine learning (ML) models, including the *SMRT* dataset (Domingo-Almenara *et al.*, 2019) for ET prediction and the *All-CCS* dataset (Zhou *et al.*, 2020) for CCS prediction. We randomly split the compounds in the *SMRT* dataset into the training and test subsets with the ratio of 9:1. For the comparison with the customized CCS prediction method ALLCCS (Zhou *et al.*, 2020), which was trained on the whole AllCCS dataset but was not shared in open-source, we used the entire ALLCCS as the training set, and used another CCS dataset from Bush Lab (Bush *et al.*, 2010) *(BushCCS*), which does not overlap with AllCCS, as the testing set for model evaluation. The sizes of the training and testing datasets for the ET and CCS prediction are summarized in Table 4.

**Table 4:** The training/test datasets and the performance of the collisional cross section (CCS) and elution time (ET) prediction. The second column shows the number of measurements and compounds in the training and testing set, respectively (separated by a comma).

| Task | # measurements<br>(# compounds) | Mean ± SD<br>Range | Methods | $R^2$ | MAE |
| --- | --- | --- | --- | --- | --- |
| CCS | 2886, 1163<br>(1721, 1151) | 170.8 ± 36.2<br>[105.9, 355.8] | AllCCS | 0.781 | 9.781 |
|  |  |  | DeepCCS | 0.850 | 9.205 |
|  |  |  | 3DMolMS | 0.957 | 6.014 |
|  |  |  | 3DMolMS-TL | 0.961 | 5.030 |
| ET | 72344, 7608<br>(72344, 7608) | 790.1 ± 206.7<br>[0.3, 1471.7] | SMRT-DLM | 0.775 | 58.534 |
|  |  |  | 3DMolMS | 0.778 | 58.459 |
|  |  |  | 3DMolMS-TL | 0.787 | 55.300 |

To compare the prediction performance, we computed the coefficient of determination (*R*^2^) and the mean absolute error (MAE) between the predicted and the experimentally measured ET and CCS. As shown in Table 4, the 3DMolMS model trained from scratch on the respective training data achieved better performance than the customized ET and CCS prediction tools SMRT-DLM (Domingo-Almenara *et al.*, 2019), ALLCCS (Zhou *et al.*, 2020), and DeepCCS (Plante *et al.*, 2019), respectively on both the *R*^2^ and the MAE metrics, while the model trained on the transferred molecular representations (denoted as ’3DMolMS-TL’) further improved the performance of 3DMolMS. In particular, the improvement in the CCS prediction is very significant, perhaps because CCS is highly correlated with the 3D structure of the compounds, which are explicitly modeled by 3DMolMS. These results suggest that the latent molecular representations learned from MS/MS spectra can be exploited to enhance the performance of predicting the chemical properties of compounds.

## 4 Conclusion

In this paper, we designed an elemental convolution-based operation to extract feature representations from 3D molecular conformations, and developed 3DMolMS, a point-based deep neural network (DNN) to predict MS/MS spectra compounds. The elemental operation is proved to be invariant to permutation and SE(3), which is essential for point-based representation learning. As our experiments showed, the 3DMolMS model can give accurate predictions of compounds’ MS/MS spectra and outperform existing MS/MS spectra prediction algorithms (e.g., NEIMS, MassFormer, and CFM-ID 4.0) as well as baseline point-based DNN models (e.g. PointNet, DGCNN, and SchNet) equipped with the same decoder as 3DMolMS for MS/MS spectra prediction. Finally, we showed that the molecular representation learned from MS/MS spectra prediction can be used to enhance the prediction of chemical properties of compounds, including the elution time (ET) and the collision cross section (CCS).

We anticipate that accurately predicted spectra can expand the reference libraries, which, combined with the predicted CCS and ET, can improve compound identification using tandem mass spectrometry. We plan to test the application of our methods to large-scale compound identification (e.g., for untargeted metabolomics) in our future works. It is also worth noting that there is plenty of room for further improvement of MS/MS spectra prediction, in particular by increasing the coverage and diversity of the compounds in the training datasets.

## Acknowledgements

We are indebted to Dr. Mingxun Wang for his invaluable advice and his effort to build the online service of 3DMolMS.

