## Supplemental Materials for "3DMolMS: Prediction of Tandem Mass Spectra from Three Dimensional Molecular Conformations"

#### Contents

**S1 Satisfaction of invariance principles** **S2**

**S2 Implementation and Training** **S3**

**S3 Experiments on Other Mass Spectra Predictors** **S3**
- S3.1 Mass Spectra Prediction Implemented from PointNet and DGCNN . . . . . . . . . . . . S3
- S3.2 Mass Spectra Prediction Implemented from SchNet . . . . . . . . . . . . . . . . . . . . . . S3
- S3.3 NEIMS and MassFormer . . . . . . . . . . . . . . . . . . . . . . . . . . . . . . . . . . . . S3

**S4 Supplementary Figures** **S4**
- S4.1 Figure S1 . . . . . . . . . . . . . . . . . . . . . . . . . . . . . . . . . . . . . . . . . . S4
- S4.2 Figure S2 . . . . . . . . . . . . . . . . . . . . . . . . . . . . . . . . . . . . . . . . . S4
- S4.3 Figure S3 . . . . . . . . . . . . . . . . . . . . . . . . . . . . . . . . . . . . . . . . . S5
- S4.4 Figure S4 . . . . . . . . . . . . . . . . . . . . . . . . . . . . . . . . . . . . . . . . . S6

#### S1 Satisfaction of invariance principles

There are two invariance principles (Qi *et al.*, 2017) that the point-based DNN should follow: the permutation invariance, and the SE(3) invariance, which guarantees the prediction of the DNN remains the same (invariant) when the $x, y, z$-coordinates in the input points are rotated and/or translated. The permutation invariance is satisfied by the main idea of point set representation because the elemental operation is symmetric on all the points. Below, we show the elemental operation 3DMolConv satisfies the SE(3) invariance. Let's restate the equations used in 3DMolConv here.

In each layer $l$, we define the 3DMolConv as:

$$x_i^{l+1} = x_i^l + \sum_{j \in \mathcal{N}(x_i^l)} d(x_i^l, x_j^l) W_1^l \circ \phi(x_i^l, x_j^l) W_2^l \circ x_j^l \tag{1}$$

The distance between two points $x_i$ and $x_j$ is computed as:

$$d(x_i, x_j) = ||x_i - x_j|| \tag{2}$$

The angle between the point vectors $x_i$ and $x_j$ encodes the information related to either the bond angle or the non-bond angles of the edge $\langle x_i, x_j \rangle$:

$$\phi(x_i, x_j) = \sum_{k \in \mathcal{N}(x_i)} e_{ij}^\top e_{ik} \tag{3}$$

To consider the representation of the whole compound, we express the representation matrix of all edges according to Eq. 3:

$$E(X) = X^\top X M = \begin{bmatrix} x_1^\top x_1 & x_1^\top x_2 & \cdots & x_1^\top x_n \\ x_2^\top x_1 & x_2^\top x_2 & \cdots & x_2^\top x_n \\ \vdots & \vdots & \ddots & \vdots \\ x_n^\top x_1 & x_n^\top x_2 & \cdots & x_n^\top x_n \end{bmatrix} M \tag{4}$$

where $M$ is a mask for $k$-nearest neighbors.

**Theorem 1.** *The edge matrix is rotation-invariant, i.e., for* $\forall R \in SO(3)$, $\forall X \in \mathbb{R}^{N \times F} (N \in \mathbb{N}^+, F \in \mathbb{N}^+)$*, i.e., it satisfies*

$$E(X) = E(XR) \tag{5}$$

*Proof.* For $\forall R \in SO(3)$, $\forall X \in \mathbb{R}^{N \times F} (N \in \mathbb{N}^+, F \in \mathbb{N}^+)$, according to the definition of edge matrix in Eq.4,

$$E(XR) = (RX)^\top (RX) M = X^\top R^\top R X M = X^\top X M \tag{6}$$

so $E(XR) = E(X)$, i.e. the edge matrix is rotation-invariant. □

Similarly, we can prove that $\phi(x_i, x_j)$ and $d(x_i, x_j)$ are both rotation-invariant. Note that the $x, y, z$-coordinates are not directly encoded after the first layer; therefore, the representations in every follow-up layer (Eq. 1) are SO(3), and the whole model is SO(3). In addition, since we always move the gravity center of $x, y, z$-coordinates in the input point set to the origin through a preprocessing step, the representations learned in the 3DMolMS model are always translation invariant. Hence, the molecular representation in 3DMolMS is SE(3) invariant.

### S2 Implementation and Training

We implemented 3DMolNet using PyTorch (Paszke *et al.*, 2017, 2019). The entire 3DMolMS model contains a total of $91,658,760$ parameters. The model is trained by Adam (Kingma and Ba, 2014) optimizer with a learning rate at 0.001 and batch size at 64. The learning rate is reduced by half when the loss does not go down in 5 epochs (implemented by `ReduceLROnPlateau` in PyTorch). We use the early stopping strategy to avoid overfitting. Specifically, the training stopped when the loss does not go down in 10 epochs. All the training and test codes of 3DMolMS are released on GitHub at `https://github.com/JosieHong/3DMolMS`, and it can also be accessed through an online service at `https://spectrumprediction.gnps2.org`.

The pre-training on multiple ion modes of Agilent QTOF spectra takes about 8 hours, and all the finetuning (i.e. finetuning on positive/negative ion mode of Agilent QTOF spectra, and finetuning on positive/negative ion mode of other QTOF spectra) takes less than 1 hour on one NVIDIA GTX 1080ti GPU. On average, the conformation generation and prediction takes about 0.22 and 0.13 seconds per compound on average.

### S3 Experiments on Other Mass Spectra Predictors

#### S3.1 Mass Spectra Prediction Implemented from PointNet and DGCNN

| Staff | PoinNet_MS | DGCNN_MS |
|---|---|---|
| embedding type | atoms point | atoms point |
| feed-forward layers type | mlp | edge conv |
| feed forward layers hidden dimensions | 64, 64, 128, 256, 512, 1024 | 64, 64, 128, 256, 512, 1024 |
| embedding dimension | 2048 | 2048 |
| decoder layers | 2048, 2048, 2048, 2048, 2048 | 2048, 2048, 2048, 2048, 2048 |
| dropout | 0.3 | 0.3 |

Table S1: Experimental settings of PointNet_MS and DGCNN_MS.

The feed-forward layers of PointNet (Qi *et al.*, 2017) are reorganized as 64, 64, 128, 256, 512, 1024, which is an advanced structure proposed in DGCNN (Wang *et al.*, 2019). The decoder for mass spectra prediction (see Fig. S1) is equipped.

#### S3.2 Mass Spectra Prediction Implemented from SchNet

We kept all the encoding settings of SchNet (Schütt *et al.*, 2018), including the three intersection block with the hidden dimension of 64, the atom-wise layers with the hidden dimension of 32, and the shifted sofpplus. Our implementation of SchNet is referred to `https://github.com/atomistic-machine-learning/schnetpack`. Also, the decoder for mass spectra prediction (see Fig. S1) is equipped.

#### S3.3 NEIMS and MassFormer

We adjusted the settings of NEIMS (Wei *et al.*, 2019) and MassFormer(Young *et al.*, 2021) as shown in Table S2, which retain most of the original settings but increase the number of the feed-forward layers from 3 to 6 enlarging the capacity of the models. There are two types of feed-forward layers: 'neims' means the same linear block proposed in NEIMS (Wei *et al.*, 2019); 'standard' means one linear layer with batch normalization and dropout.

The training process is changed to pretraining on multiple precursor types and fine-tuning on a certain precursor type, which is the same as the training process of other predictors. All the implementations of NEIMS and MassFormer are available at `https://github.com/Roestlab/massformer`.

| Staff | NEIMS | MassFormer |
|---|---|---|
| embedding type | fingerprint | molecular graph |
| feed-forward layers type | neims | standard |
| feed-forward layers hidden dimension | 1000 | 1000 |
| feed-forward layers number | 6 | 6 |
| feed-forward layers skip | False | True |
| gate prediction | True | True |
| dropout | 0.15 | 0.15 |
| output normalization | l1 | l1 |
| model seed | 6666 | 6666 |

Table S2: Experimental settings of NEIMS and MassFormer.

### S4 Supplementary Figures

#### S4.1 Figure S1

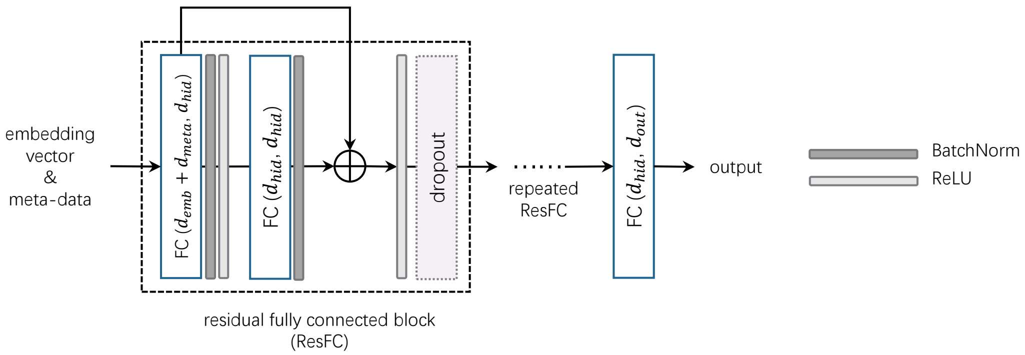


Figure S1: Illustration of mass spectra decoder. The fully connected block is repeated five times in all experiments.

#### S4.2 Figure S2

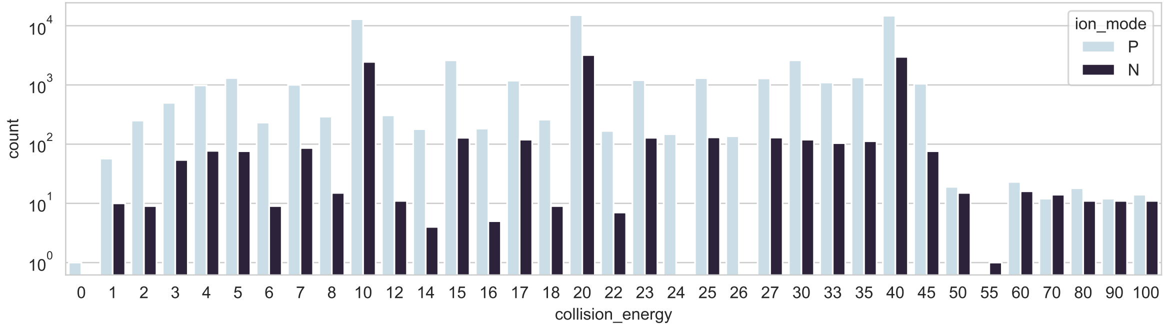


Figure S2: Distribution of collision energies in the Agilent QTOF spectra from Agilent PCDL and NIST20.

#### S4.3 Figure S3

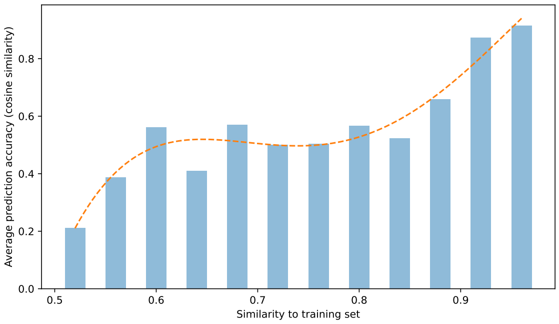


(a) 40 eV, $[\mathrm{M}+\mathrm{H}]^+$

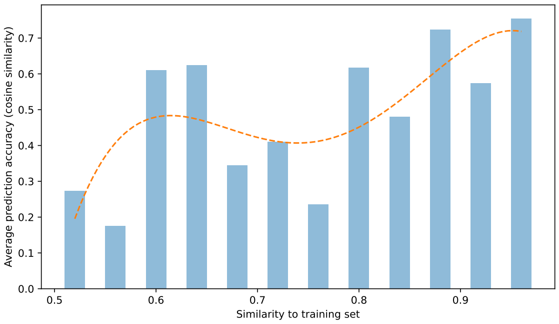


(b) 40 eV, $[\mathrm{M}-\mathrm{H}]^-$

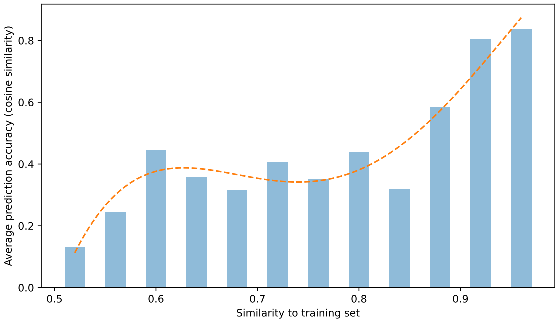


(c) 20 eV, $[\mathrm{M}+\mathrm{H}]^+$

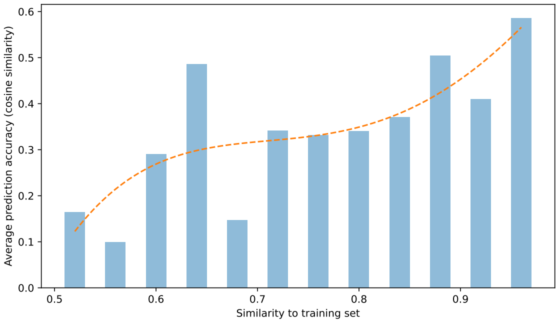


(d) 20 eV, $[\mathrm{M}-\mathrm{H}]^-$

Figure S3: The accuracy of predicted spectra is correlated with the max Bulk Tanimoto similarity between the test molecule and the training molecules.

#### S4.4 Figure S4

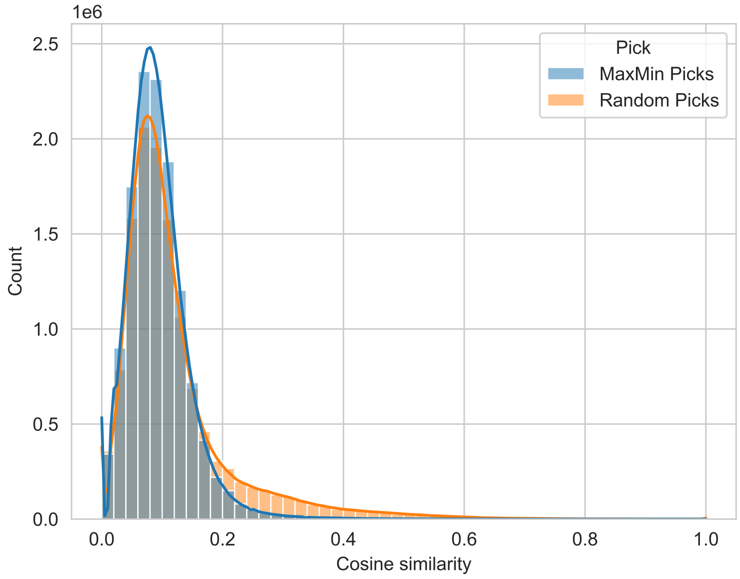


(a) The training set using MaxMin Pick (Ashton *et al.*, 2002) has lower cosine similarities (i.e. higher diversity) than the training set using random picking.

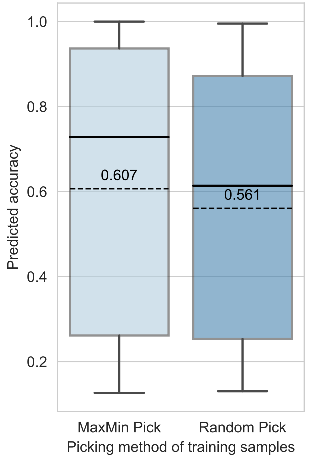


(b) 3DMolMS trained in diversity training samples (i.e. training samples picked by MaxMin Pick) achieves higher predicted accuracy than using less diversity training samples.

Figure S4: Experiments of using different diversity training samples.
